# Hantavirus stability and inactivation

**DOI:** 10.1101/2025.11.20.689437

**Authors:** Léna Vandenabeele, Abraham Ayanwale, Thomas Pietschmann, Benjamin E. Nilsson-Payant

## Abstract

Hantaviruses are zoonotic viruses that can cause highly pathogenic disease, including hantavirus cardiopulmonary syndrome (HCPS) and haemorrhagic fever with renal syndrome (HFRS), in humans with case-fatality rates of up to 50%. However, our understanding of the basic viral life cycle and the underlying causes of viral pathogenesis remains sparse, in large part due to a lack of molecular biology tools for hantaviruses and the need to work in high-containment laboratory facilities with these viruses. Here, we investigated the kinetics of infectious Tula virus (TULV) particle production in Vero E6 cells and subsequent stability in cell culture media. In addition, we evaluated the stability of infectious virus particles in response to different physical and environmental stresses, including heat, freezing, dehydration and UV exposure, answering key questions about the environmental transmission potential of hantaviruses. Interestingly, we observed a remarkable stability of TULV when stored at room temperature or colder, as well as after dehydration, which suggests that hantaviruses could remain infectious for a sustained period of time after being secreted by their host species. Subsequently, we determined the ability of commonly used virus inactivation methods, including RNA and protein extraction buffers, to inactivate TULV both in a cell-free and cell-associated context and found that TULV was efficiently inactivated by all these methods similar to other enveloped RNA viruses. Finally, we successfully validated the complete inactivation using these inactivation methods using the highly pathogenic HCPS-causing New World Andes virus (ANDV) and the HFRS-causing Old World Hantaan virus (HTNV). These results provide valuable information about safe and effective inactivation methods of viral samples and about the environmental risk potential of hantaviruses.

**Author summary:** Hantaviruses are ubiquitous rodent viruses and contain some of the most lethal known zoonotic viruses, including Andes virus (ANDV) and Hantaan virus (HTNV), with no FDA-or EMA-approved antiviral treatment or prevention options available. However, studying the molecular biology and pathogenesis of these viruses is significantly impeded by the need for biosafety level (BSL) 3 (or higher) containment facilities for most hantaviruses and a general lack of molecular biology tools. Our study provides a comprehensive analysis of the stability of infectious hantavirus particles in response to different physical and chemical stresses. We demonstrate that a diverse range of hantaviruses are effectively and quickly inactivated by commonly used generic viral inactivation methods. However, we also provide evidence for an inherent stability under environmental conditions that could enable prolonged transmission potential of hantaviruses, even after secretion from their host species. These data are essential information to design safe inactivation methods of infectious hantavirus material and offers insights into the environmental risks of hantavirus infections with implications for laboratory safety and public health measures.

## Background

Orthohantaviruses (henceforth called hantaviruses) are a genus of segmented negative-sense RNA viruses within the *Hantaviridae* family that contain some of the most lethal known human pathogens. Ordinarily, hantaviruses circulate in their natural hosts – primarily rodents, but also shrews, moles, bats and other small mammals – where they are not thought to cause any disease [1,2]. However, some hantaviruses are known to be able to zoonotically infect humans and cause serious illnesses. Some Old World hantaviruses – including Hantaan virus (HTNV) and Seoul virus (SEOV) – can cause haemorrhagic fever with real syndrome (HFRS) in humans, which in severe cases is characterized by hypotension, vascular leakage, acute shock, acute kidney failure and case-fatality rates of up to 15% [3]. In Europe, Puumala virus (PUUV) can cause a mild version of HFRS called Nephropathia epidemica (NE), which is associated with fever, gastrointestinal symptoms and impaired renal function and case-fatality rates of less than 1% [4]. New World hantaviruses – including Andes virus (ANDV) and Sin Nombre virus (SNV) – are known to cause a severe respiratory disease called hantavirus cardiopulmonary syndrome (HCPS), which is characterized by initial flu-like symptoms which can rapidly progress to severe respiratory distress, pulmonary vascular leakage and respiratory failure with mortality rates of up to 40% [5].

Natural host reservoir species shed and transmit viral particles in their excreta, *e.g.* their saliva, urine and faeces and zoonotic infection of humans primarily occurs through the inhalation of these excreta, *e.g.* in the form of aerosolized dust particles [1]. Humans usually are dead-end hosts for hantaviruses, but human-to-human transmission, including nosocomial infections, has been observed for ANDV, causing several local epidemics in Argentina and Chile [6–8].

While it is impossible to accurately estimate the burden of human hantavirus infections due to their often-asymptomatic nature or very generic flu-like symptoms in mild cases, it has been estimated that globally 60,000-100,000 annual cases are recorded every year, with the majority of cases caused by HTNV and SEOV in China [1]. While hantavirus cases have reduced over the past decades in China, over 1.5 million cases of HFRS were recorded between 1950 and 2007, with a peak of 115,804 cases in 1986 [9,10]. In Europe, PUUV is the main cause of human infection accounting for usually less than 10,000 confirmed cases of human infections, with Finland, but also Sweden and Germany reporting the majority of cases [4]. In the Americas, human infection with New World hantaviruses occur much more sporadic and only a few thousand cases in total (the majority in Argentina and Chile) have been recorded since the discovery of new World hantaviruses in 1993 [1,11]. It is important to note however, that the actual case numbers for all hantaviruses are likely to be significantly higher.

While efforts are ongoing to identify and develop specific antiviral treatment options and vaccines – including clinical trials using DNA vaccines and broadly neutralizing monoclonal antibodies, there are currently no FDA- or EMA-approved antiviral treatments or vaccines available [12]. In consequence of this and taking their severe pathogenic potential into account, most hantaviruses need to be handled in high containment facilities. This significantly hampers the ease of basic and translational research, as the virus can only be handled in select laboratory facilities and requires stringent inactivation methods in order to be retrieved from the high-containment facilities. It is therefore important to gain a better understanding of how infectious material safely and effectively can be inactivated and how durable infectious particles remain under different environmental stresses. In this study, we utilized Tula virus (TULV) – a mostly apathogenic close relative of PUUV – to evaluate its ability to retain its infectious capacity in response to different environmental and physical stresses, including exposure to heat, UV irradiation, long-term exposure at different temperatures, freezing and dehydration. Furthermore, we sought to determine whether different inactivation and lysis methods commonly used in lab protocols are able to effectively inactivate TULV both in a cellular and cell-free context. Finally, we validated whether our findings for TULV could be cross-applied to highly pathogenic HFRS-causing (HTNV) and HCPS-causing (ANDV) viruses.

## Methods

### Cell and virus culture

African green monkey kidney epithelial cells (Vero E6; ATCC, CRL-1586) were commercially obtained. Cells were maintained in Dulbecco’s Modified Eagle Medium (DMEM) supplemented with 10% foetal bovine serum (FBS), L-Glutamine, MEM Non-Essential Amino Acids and 100 U/ml penicillin and 100 μg/ml streptomycin and cultured at 37°C and 5% CO_2_.

Tula virus (TULV; *Orthohantavirus tulaense*) strain Moravia/Ma5302V/94 was a kind gift by Rainer Ulrich (Friedrich-Loeffler-Institut, Germany). Tula/ Moravia/Ma5302V/94 virus was originally isolated from the lungs of infected *Microtus arvalis* trapped in Tvrdonice, South Moravia (Czech Republic) by passaging once in a hantavirus-free laboratory *Microtus arvalis* animal and subsequently in Vero E6 cells [13,14]. Andes virus (ANDV; *Orthohantavirus andasense*) strain Chile-9717869 was a kind gift by Piet Maes (KU Leuven, Belgium). Andes/Chile-9717869 virus was originally isolated from the kidneys and lungs of an infected wild *Oligoryzomys longicaudatus* trapped during an outbreak of HCPS in Chile in 1997 by passaging in Vero E6 cells [15,16]. Hantaan virus (HTNV; *Orthohantavirus hantanense*) strain 76-118 was acquired from the European Virus Archive Global (EVAg) repository (Ref-SKU: 008V-EVA1471) through its partner institution BMC-SAS (Bratislava, Slovakia). Hantaan/76-118 virus was originally isolated from the lungs of an infected wild *Apodemus agrarius* in Songnaeri, North Korea, in 1976 and subsequently passaged in A549 cells [17,18].

All viruses were propagated in Vero E6 cells in DMEM supplemented with 2% FBS, L-Glutamine, MEM Non-Essential Amino Acids, 100 U/ml penicillin and 100 μg/ml streptomycin. Virus-containing cell culture supernatants were repeatedly collected between 5 to 14 days post-infection (dpi) and replaced with fresh infection media. Pooled viral stocks were purified and concentrated using Amicon Ultra-15 centrifugal filter units (100-kDa molecular weight cut-off). Infectious virus titres were determined by immuno-plaque assays as described below.

All work involving infectious ANDV and HTNV was performed in the biosafety level 3 (BSL-3) facility of the Hannover Medical School (MHH) in accordance with its institutional biosafety requirements.

### Immuno-plaque assay

Infectious viral titres were determined by immuno-plaque assays as previously described in Vero E6 cells [19]. Vero E6 cells were infected with sequential 10-fold dilutions of virus and overlayed with 1.2% Avicel CL-611 in MEM containing 2% FBS, 10 mM HEPES, MEM Non-Essential Amino Acids and 100 U/ml penicillin and 100 μg/ml streptomycin. At 7 days post-infection, cells were fixed for 30 min at room temperature in 3.7% formaldehyde and permeabilized for 30 min at room temperature in 0.5% Triton X-100. Infected cells were immunostained for 2 h at room temperature with anti-NP monoclonal antibody clones TULV1 [20] for TULV and ANDV (a kind gift by Rainer Ulrich, Friedrich-Loeffler-Institut) or B5D9 for HTNV (Progen, cat# B5D9-C) and visualized with an IRDye 800CW-conjugated anti-mouse IgG (LI-COR, cat# 926-32210) secondary antibody. Fluorescent signal of stained plaques was detected using an Odyssey CLx imaging system (LI-COR) and analysed with the Image Studio software (LI-COR).

### Viral propagation assay

The continuous production of infectious TULV particles in Vero E6 cells was measured over time as following. Vero E6 cells were infected in triplicates with TULV at an MOI of 0.001 in infection medium (DMEM supplemented with 2% FBS, L-Glutamine, MEM Non-Essential Amino Acids, 100 U/ml penicillin and 100 μg/ml streptomycin). At 1 h post-infection (hpi), the viral inoculum was removed and replaced with fresh infection medium. At the indicated time points either 5% or 100% of cell culture supernatant was collected and replaced with fresh infection medium. Collected viral samples were frozen and stored at -80°C before all infectious titres were determined by immuno-plaque assays as described above.

### Physical inactivation of cell-free viral particles

To determine the effects of dehydration on viral particles, approximately 1 × 10^6^ PFU of virus were placed in a sterile cell culture plate and left to air dry in a biosafety cabinet for approximately 2 h. After complete dehydration, dried viral solutions were reconstituted in 300 µl DMEM supplemented with 2% FBS, L-Glutamine, MEM Non-Essential Amino Acids, 100 U/ml penicillin and 100 μg/ml streptomycin, incubated at room temperature for 5 min and mixed extensively by pipetting. As a control approximately 1 × 10^6^ PFU of virus were placed in a closed microcentrifuge tube and exposed to the same temperatures for the same time but not allowed to air-dry. To determine the effects of freezing viral solutions, freshly thawed aliquots of virus were flash-frozen on dry ice before being rethawed. For thermal inactivation of viral particles, approximately 1 × 10^6^ PFU of virus were placed in PCR tubes and exposed to the indicated temperatures for the indicated time in a preheated thermocycler. To determine thermal stability of viral particles, approximately 1 × 10^6^ PFU of virus were stored in the dark at 4°C, 21°C or 37°C. At the indicated time points, aliquots were frozen and stored at -80°C. For ultraviolet (UV) irradiation of viral particles, approximately 1 × 10^6^ PFU of virus were placed in a sterile cell culture dish and exposed to indicated amounts of short wavelength UV-C irradiation (254 nm) in a CX-2000 UV Crosslinker (UVP). The cell culture dishes were placed approximately 10 cm below the UV-C light source. Infectious viral titres before and after all treatments were determined in parallel by immuno-plaque assay as described above.

### Chemical inactivation of cell-free viral particles

For chemical inactivation of cell-free viral particles, approximately 1 × 10^6^ PFU of virus were mixed at a 1:1 vol/vol ratio for 10 min at room temperature with the chemicals at the final concentrations indicated in Table 1. PBS was used as a non-inactivating control. Commercial RNA extraction buffers were used at the indicated concentrations according to the manufacturers’ instructions. Subsequently, solutions were diluted in 10 ml PBS in order to reduce the concentration of cytotoxic chemicals and viral particles were concentrated using Amicon Ultra-15 Centrifugal Filter Units (100-kDa molecular weight cut-off). Retained concentrates (approximately 0.2 ml) were diluted once more in 10 ml PBS for an approximately 2,500-fold final dilution of inactivating chemicals and reconcentrated using the same Amicon Ultra-15 Centrifugal Filter Units. Retained solutions were titrated for infectious viral particles using immuno-plaque assays as described above.

**Table 1:** Inactivation methods for cell-free infectious particles.

| ***Chemical*** | ***Final concentration*** | ***Exposure time*** |
| --- | --- | --- |
| Ethanol | 20%, 40%, 70% | 10 min |
| Formaldehyde | 1%, 4% | 10 min |
| Sodium dodecyl sulfate (SDS) | 1% | 10 min |
| Nonidet P-40 substitute (NP-40) | 0.5% | 10 min |
| Triton X-100 | 0.1%, 1% | 10 min |
| TRI Reagent | 3:1 (75%) | 10 min |
| Viral RNA Buffer<br>(Zymo Research, cat# R1034-1) | 2:1 (66.67%) | 10 min |
| Buffer AVL (QIAGEN, cat# 19073) | 4:1 (80%) | 10 min |
| Binding Buffer (Roche, cat# 11858882001) | 2:1 (66.67%) | 10 min |

### Chemical inactivation of intracellular viral material

For chemical inactivation of intracellular viral particles, approximately 240,000 Vero E6 cells were infected at an MOI of 1 (TULV) or 0.2 (HTNV) in DMEM supplemented with 2% FBS, L-Glutamine, MEM Non-Essential Amino Acids, 100 U/ml penicillin and 100 μg/ml streptomycin. At 2 dpi, cell culture supernatants were removed off infected cell monolayers and replaced with 100 μl of the indicated chemicals (Table 2) or PBS as control and frozen at -80°C to facilitate full cellular rupture. After thawing of cells, 50% of cell lysates were used to infect approximately 240,000 naïve Vero E6 cells in triplicates in 1 ml DMEM supplemented with 2% FBS, L-Glutamine, MEM Non-Essential Amino Acids, 100 U/ml penicillin and 100 μg/ml streptomycin (resulting in an approximately 20-fold dilution of chemicals). After 9 to 20 dpi, the concentration of infectious viral particles in the cell culture supernatants were determined using immuno-plaque assays as described above.

**Table 2:**
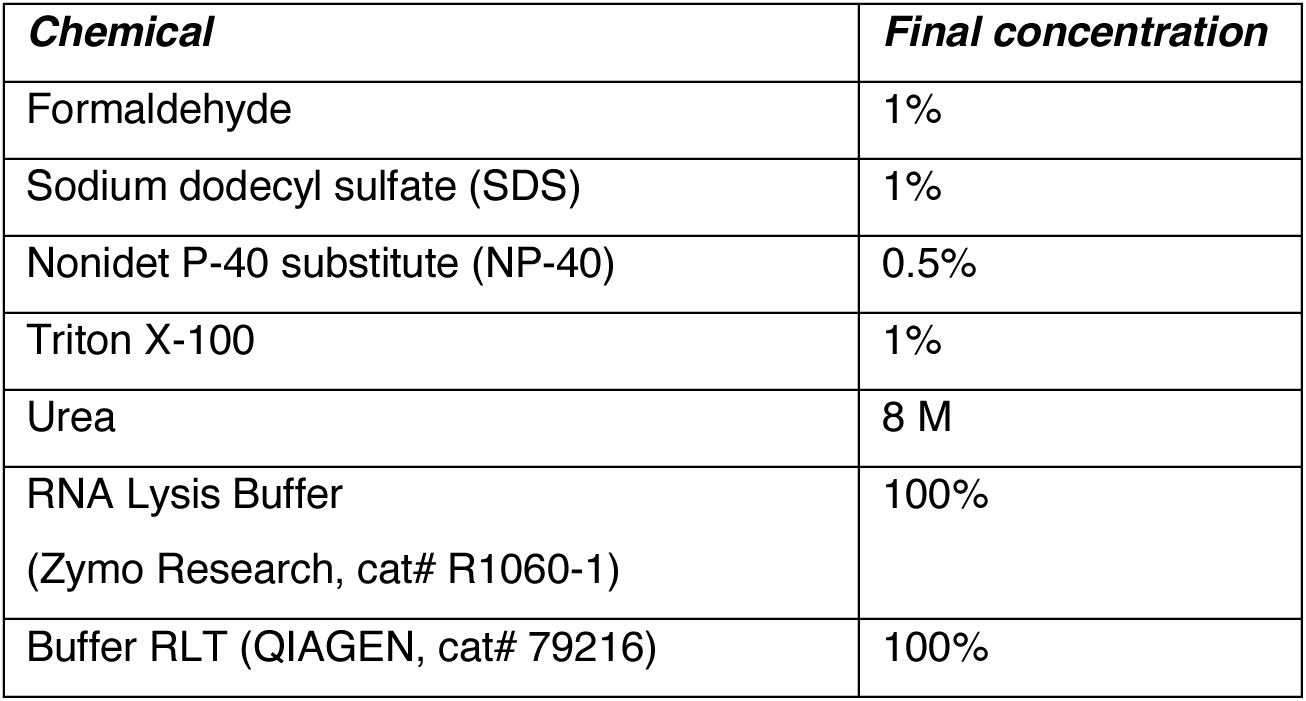
Inactivation methods for intracellular infectious material.

| ***Chemical*** | ***Final concentration*** |
| --- | --- |
| Formaldehyde | 1% |
| Sodium dodecyl sulfate (SDS) | 1% |
| Nonidet P-40 substitute (NP-40) | 0.5% |
| Triton X-100 | 1% |
| Urea | 8 M |
| RNA Lysis Buffer<br>(Zymo Research, cat# R1060-1) | 100% |
| Buffer RLT (QIAGEN, cat# 79216) | 100% |

## Results

### TULV virion production and stability in cell culture media

Hantaviruses are relatively unique within RNA viruses in that they do not cause any significant viral-induced cytopathic effects (CPE). It has previously been shown that cells can be constitutively infected with hantaviruses and continuously produce and release viral particles into the cell culture media [21,22,20]. In order to get a better insight into the kinetics of infectious virion production and their subsequent stability in cell culture media, we infected Vero E6 cells with the mostly apathogenic TULV and measured the concentration of both nascently produced and cumulatively produced infectious viral particles in the cell culture supernatant over three weeks of infections (Fig 1A). We observed no infectious viral particles at 1 and 3 dpi, with low titres after 5 dpi and then a rapid increase in virion release and a clear peak in infectious viral particle production around 10 dpi, which then subsequently decreased again and plateaued between 14 and 21 dpi (Fig 1B). Notably, the cumulative and nascent infectious titres remained essentially identical to each other at each collection time. This indicates a short half-life of infectious viral particles in cell culture media at 37°C, meaning that the measured cumulative titres only reflect nascently produced viral particles and that older particles quickly lose their infectious potential.

**Figure 1:**
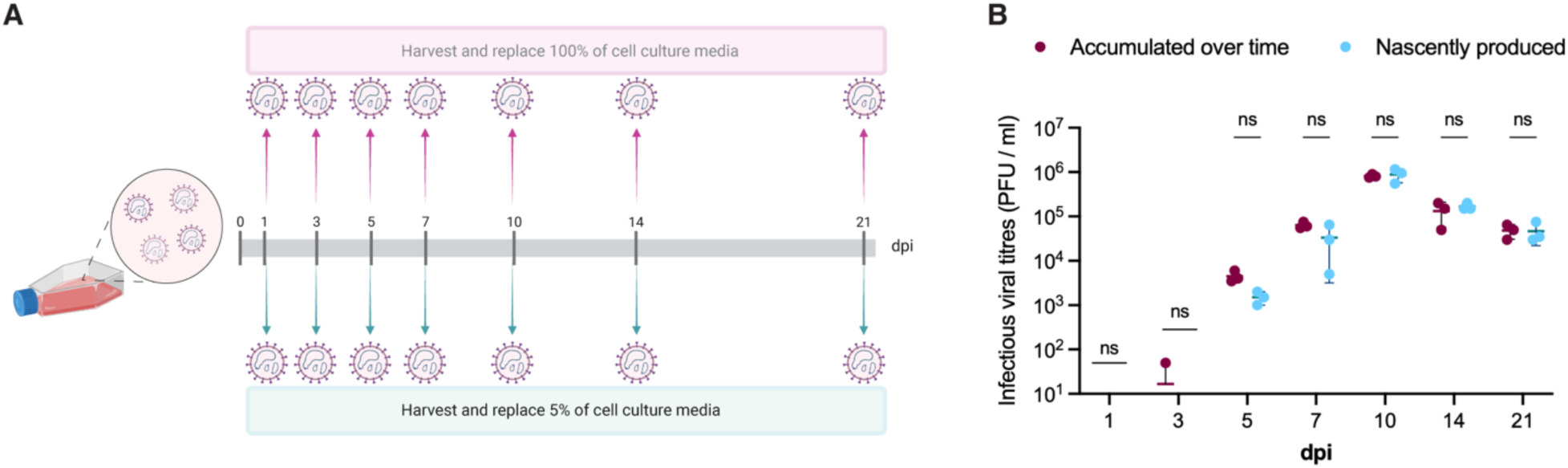
Infectious TULV particle production in Vero E6 cells. (A) Schematic of the experimental setup: Vero E6 cells were infected with TULV (MOI = 0.001) and either 100% or 5% of cell culture media were collected and replaced at the indicated time points before determining infectious viral titres. (B) Infectious viral titres of TULV at the indicated time points as determined by immuno-plaque assay. Individual dots represent independent biological replicates (n = 3). Statistical significance was determined by two-way ANOVA followed by Šidák’s correction for multiple comparisons: ns, p > 0.05.

### Stability and inactivation of TULV under physical stress

The ability of different viruses to survive different physical and environmental stresses and maintain infectivity differs vastly. Here, we tested the ability of TULV to withstand environmental stresses, including temperature, freezing, dehydration and UV exposure.

Ultraviolet (UV) irradiation is generally known to be able to inactivate RNA viruses, although the efficacy for different viruses varies depending on the wavelength and exposure time. Here we first exposed TULV in cell culture media to UV irradiation at 254 nm, resulting in a significant reduction (∼10-fold) in infectious viral titres at 300 mJ/cm^2^ UV exposure, with titres dropping even further at higher doses (Fig 2A). Next, we subjected TULV to a single freeze-thaw cycle and determined infectious viral titres before and after freezing. Infectious titres were slightly, but not significantly reduced, suggesting that TULV might be well equipped to tolerate freezing temperatures in the environment (Fig 2B).

**Figure 2:**
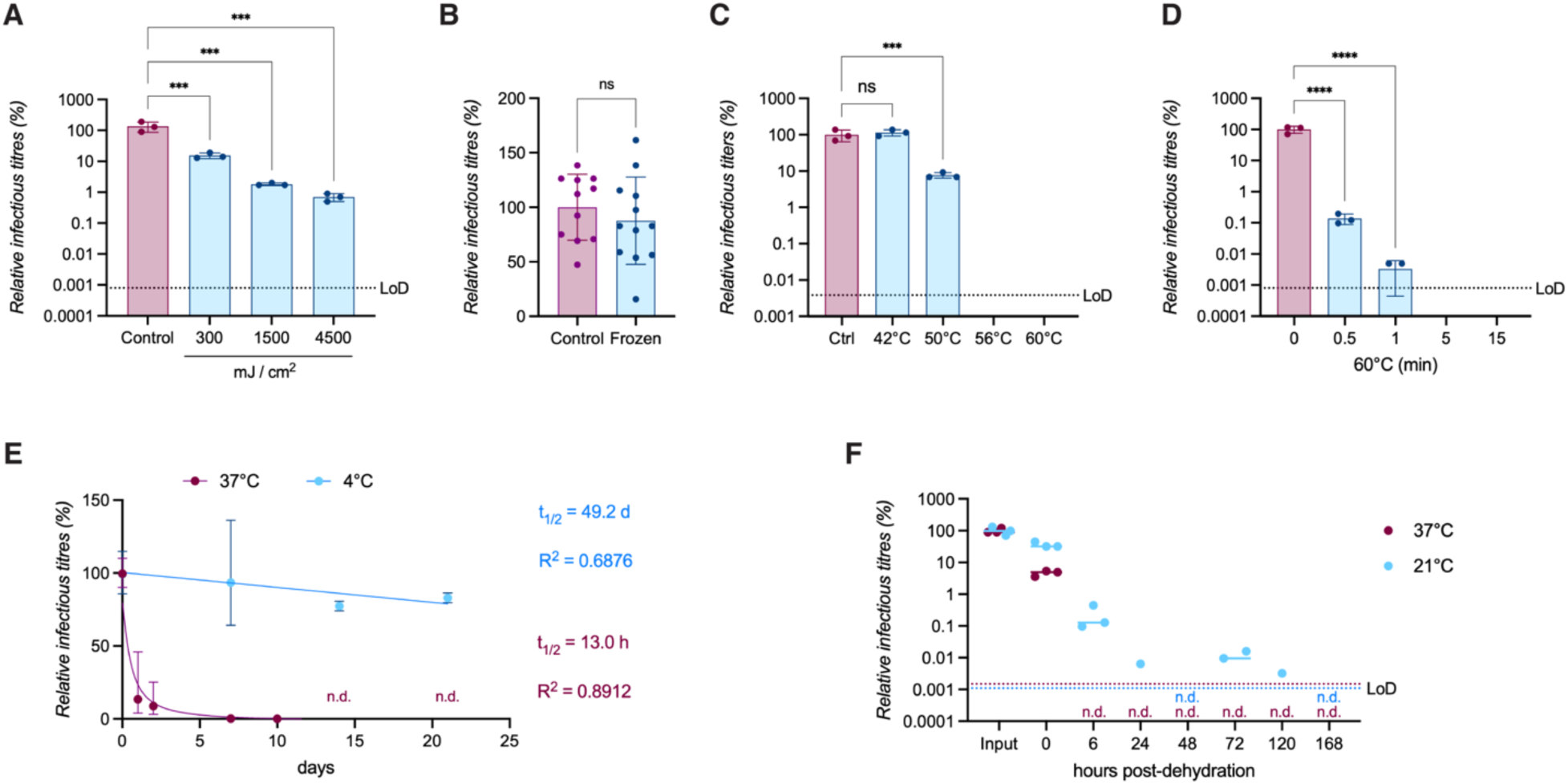
Environmental and physical stability of TULV particles. (A) Relative TULV infectious titres before and after UV (λ = 254 nm) irradiation. (B) Relative TULV titres before and after flash-freezing of virus stocks. (C) Relative TULV infectious titres before and after 15 min exposure to the indicated temperatures. (D) Relative TULV infectious titres before and after incubation at 60°C for the indicated times. (E) Relative TULV infectious titres before and after incubation at 37°C or 4°C for the indicated times (n.d. = not detected). Simple linear (for 4°C data) or nonlinear (for 37°C data) regression was used to determine the half-life (t_1/2_) of infectious virus particles stored at 4°C (blue) or 37°C (red). (F) Relative TULV infectious titres before and after dehydration at room temperature. All graphs represent relative data from immuno-plaque assays normalised to the control condition, with dots depicting independent biological replicates and error bars the standard deviation of the mean. The dotted lines represent the limit of detection (LoD) for the relevant graph. Statistical significance was determined by one-way ANOVA followed by Dunnett’s correction for multiple comparisons: **** (p > 0.0001), *** (p > 0.001), ** (p > 0.01), * (p > 0.05), ns (p > 0.05).

Enveloped RNA viruses are generally known to be efficiently inactivated by heat denaturation. We therefore sought to determine how stable TULV remains at elevated temperatures and how quickly it is inactivated. After 15 min of exposure at 42°C, no significant loss of viral titre was detected. However, at 50°C less than 10% of infectious viral particles remained and at 56°C or higher a complete loss of infectious viral titres (>26,000-fold reduction over the limit of detection) was recorded (Fig 2C). In order to ascertain how quickly this loss of viral titre manifests itself, we measured inactivation at 60°C over time and observed an immediate significant loss of viral titres after 30 seconds (∼700-fold reduction) and near complete (∼30,000-fold) inactivation after 1min (Fig 2D). At 5 or 15 min no infectious virus could be detected (>369,000-fold reduction over the limit of detection).

Due to its proven thermolability, we next sought to determine for how long TULV particles retain their infectivity when stored at different temperatures. We stored virus aliquots in the dark at 4°C or at 37°C and transferred viral samples after the indicated storage periods to -80°C and measured viral titres of all samples in parallel. At 37°C, we observed an immediate significant loss of infectious viral titres after 24 h (∼3.5-fold), which continued until infectious virus was last detected at 10 days (∼6,000-fold reduction) and a complete loss of detectable infectious viral particles between 10 and 14 days, with an estimated half-life of approximately 13 h (Fig 2E). At 4°C, we could only detect an ∼20% reduction in infectious viral titres for the entire measured period (21 days), with an estimated half-life of approximately 49 days.

As viral particles rarely exist in solution for prolonged times in the environment, it is important to test the effect of dehydration on viral infectivity. We therefore air-dried TULV at room temperature on a sterile plastic surface inside a cell culture plate until virus suspensions have been fully dehydrated. At this point dehydrated virus samples were stored in the dark at either room temperature (21°C) or at 37°C, before viral particles were rehydrated using cell culture media and infectious virus titres were measured. The act of dehydration itself resulted in significant loss of infectious virus titres with only 5-36% of viral particles remaining infectious compared to before hydration (Fig 2F). When subsequently stored at 37°C, no infectious virus could be detected after 6 h. Storage at room temperature resulted in a rapid loss of virus titres, but low levels of virus just about the limit of detection could be sporadically detected until 5 days, with a complete less of infectious viral titres after 7 days (∼90,000-fold reduction over the limit of detection).

Overall, these data suggest, that while TULV can easily be inactivated at high (≥56°C) temperatures or by UV irradiation, it is able to retain significant infectivity at low temperatures or after freezing, potentially aiding its ability to survive outside of hosts in the environment for long periods of time. In addition, even though dehydration of virus particles has a profound effect on viral infectivity, it could still be detected after several days at ambient temperatures.

### Chemical inactivation of extracellular viral particles

We demonstrated above that TULV remains infectious on surfaces and suspended in liquids for considerable amounts of time and thus can pose a hazard to humans (Fig 2). In addition, many experimental techniques involve handling of liquids containing infectious virus. It is therefore important to gain a better understanding of how cell-free infectious material can effectively and safely be inactivated by chemical agents for further processing at lower biosafety conditions.

Therefore, we tested the ability of different chemical compounds commonly used in disinfectants and inactivation buffers to effectively reduce infectious viral titres in isolation. In brief, we incubated approximately 1×10^6^ plaque forming units (PFU) of TULV in a solution with the indicated reagents at the indicated final concentrations as detailed in Table 1. Subsequently we performed a buffer exchange of virus samples in order to reduce or eliminate potentially cytotoxic concentrations of chemical agents. Thusly purified virus solutions were titrated to infectious viral particle concentrations.

We observed that incubation of virus in solution with ethanol at a final concentration of 40% or higher resulted in complete (≥100,000-fold reduction) inactivation of virus infectivity, whereas a final concentration of 20% ethanol only reduced viral titres 17.8-fold (Fig 3A). Similarly, exposure of virus to formaldehyde at 1% or higher final concentration also resulted in complete (≥100,000-fold reduction) inactivation of viral particles (Fig 3B). Many experimental methods depend on the extraction of protein or nucleic acid contents for further analysis. We therefore tested the ability of typical lysis or extraction buffers for both protein- or RNA-based methods to fully inactivate TULV in a cell-free context. For protein extraction buffers we tested commonly used ionic (*e.g.* SDS) and non-ionic (*e.g.* NP-40 and Triton X-100) detergents that are commonly used to disrupt lipid cell membranes and lyse cells (Fig 3C). Out data showed that exposure to 0.5% NP-50 and 1% Triton X-100 fully inactivated (≥100,000-fold reduction in infectious viral titres) cell-free viral solutions, whereas exposure to 1% SDS (∼5,700-fold reduction) or 0.1% Triton X-100 (∼450-fold) significantly, yet incompletely inactivated TULV in the given time frame (Fig 3C). In addition, we tested commonly used commercial RNA extraction buffers, including TRI Reagent (Zymo Research), Viral RNA Buffer (Zymo Research), AVL Buffer (QIAGEN) and High Pure Binding Buffer (Roche) according to the manufacturers’ instructions (Fig 3D). Here, we could demonstrate full inactivation of viral particles (≥100,000-fold reduction in infectious viral titres).

**Figure 3:**
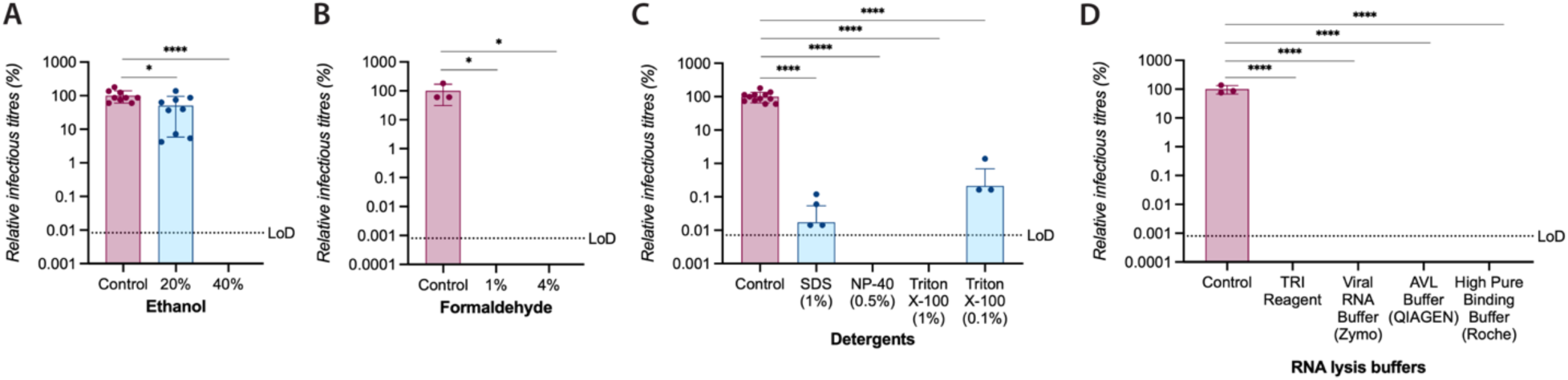
Chemical inactivation of cell-free TULV particles. Relative TULV titres before and after treatment with (A) ethanol, (B) formaldehyde, (C) detergents and (D) RNA lysis buffers. All graphs represent relative data from immuno-plaque assays normalised to the control condition, with dots depicting independent biological replicates and error bars the standard deviation of the mean. The dotted lines represent the limit of detection (LoD) for each graph. Statistical significance was determined by one-way ANOVA followed by Dunnett’s correction for multiple comparisons: **** (p > 0.0001), *** (p > 0.001), ** (p > 0.01), * (p > 0.05), ns (p > 0.05).

These data demonstrate the complete and efficient inactivation of TULV particles in solution using standard chemical inactivation methods.

### Chemical inactivation of infected cells

In Figure 3 we demonstrated that cell-free viral particles could efficiently be inactivated using commonly used buffers for cell fixation or RNA and protein extraction. Here, we tested the ability of different chemical compounds to inactivate TULV infected cells. In brief, we infected Vero E6 cells with TULV at an MOI of 1 and at 48 hpi removed cell culture supernatants and treated cells with the indicated chemicals (or PBS as a non-inactivating control) followed by freezing of cells to facilitate rupturing of cell membranes. Cellular lysates were subsequently transferred onto naïve Vero E6 cells and incubated for 9 days before measuring whether infection of naïve cells was possible and infectious viral particles could be detected in cell culture supernatants. Since we demonstrated previously that the half-life of TULV at 37°C is extremely low, we are not comparing differences in measured viral titres but rather look for the detection of any virus as a measure for whether cellular inactivation was completely successful or not.

We found that fixation of cells with ≥1% formaldehyde fully inactivated TULV infected cells and prevented infection of naïve cells (Fig 4A). When using detergents commonly used in protein extraction buffers, we could not detect any infectious virus in the supernatant after inactivation with 0.5% NP-40 or 1% Triton X-100, however, lysis of cells in 8 M urea, 1% SDS or 0.1% Triton X-100 alone was, albeit significantly reducing titres, not sufficient to fully inactivate TULV infected cells (Fig 4B). Cell lysis with commercial RNA extraction buffers, including TRI Reagent (Zymo Research), RNA Lysis Buffer (Zymo Research) and RLT Buffer (QIAGEN), led to a complete loss of detectable infectious viral titres (Fig 4C). Together, these data complement our previous findings and establish that intracellular infectious material also can be completely and efficiently be inactivated using commonly used fixation methods and lysis buffers.

**Figure 4:**
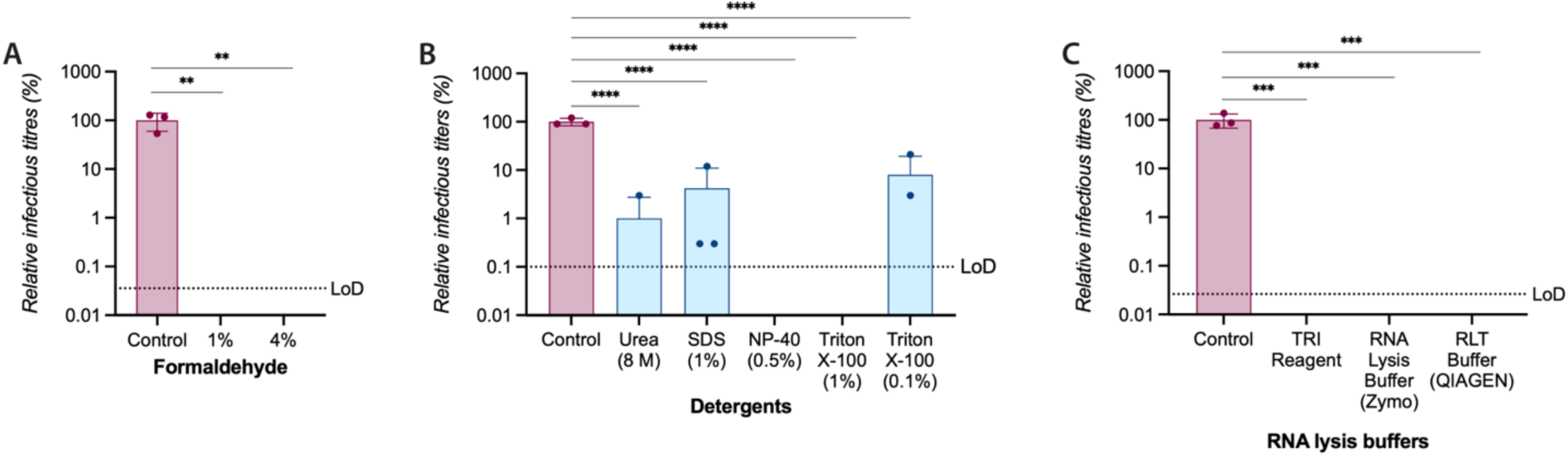
Chemical inactivation of TULV infected cells. Vero E6 cells were infected with TULV (MOI = 1) and 2 dpi, cells were inactivated using (A) formaldehyde, (B) detergents or (C) RNA lysis buffers or a PBS control and ruptured by freeze-thawing. Cell lysates were used to infect naïve Vero E6 cells and infectious titres were determined at 9 dpi. All graphs represent relative data from immuno-plaque assays normalised to the control condition, with dots depicting independent biological replicates (n = 3) and error bars the standard deviation of the mean. The dotted lines represent the limit of detection (LoD) for each graph. Statistical significance was determined by one-way ANOVA followed by Dunnett’s correction for multiple comparisons: **** (p > 0.0001), *** (p > 0.001), ** (p > 0.01), * (p > 0.05), ns (p > 0.05).

### Validation of inactivation methods using highly pathogenic hantaviruses

TULV is thought to be mostly apathogenic and therefore can safely be handled under biosafety level 2 conditions. Due to the genetic and structural similarities within the *Orthohantavirus* genus of the *Hantaviridae* family, it is a reasonable assumption that other members within this group – including highly pathogenic hantaviruses – will respond very similarly to these inactivation methods. We therefore tested whether Andes virus (ANDV), the prototypic HCPS-causing New World hantavirus, and Hantaan virus (HTNV), the prototypic HFRS-causing Old World hantavirus, displayed similar stability and could be efficiently and sufficiently inactivated by both high temperatures and chemical inactivation methods, which would allow inactivated samples to be transferred out of the biosafety level 3 facility. Initially, we tested whether both ANDV and HTNV exhibited similar stability under physical stresses as TULV. First, we dehydrated ANDV and HTNV at room temperature and measured infectious viral titres before and after dehydration (Fig 5A). We observed that for both viruses a similar proportion of infectious viral particles remained immediately after dehydration (∼35%) as for TULV (Fig 2F). Next, we compared the effects of a single freeze-thaw cycle on virus stocks (Fig 5B). Here, we observed for HTNV a non-significant reduction in infectious titres, whereas ANDV titres dropped significantly (∼37%). Finally, we validated that similar to TULV heat exposure at 60°C for 5 min was sufficient to fully inactivate both ANDV and HTNV (Fig 5C).

**Figure 5:**
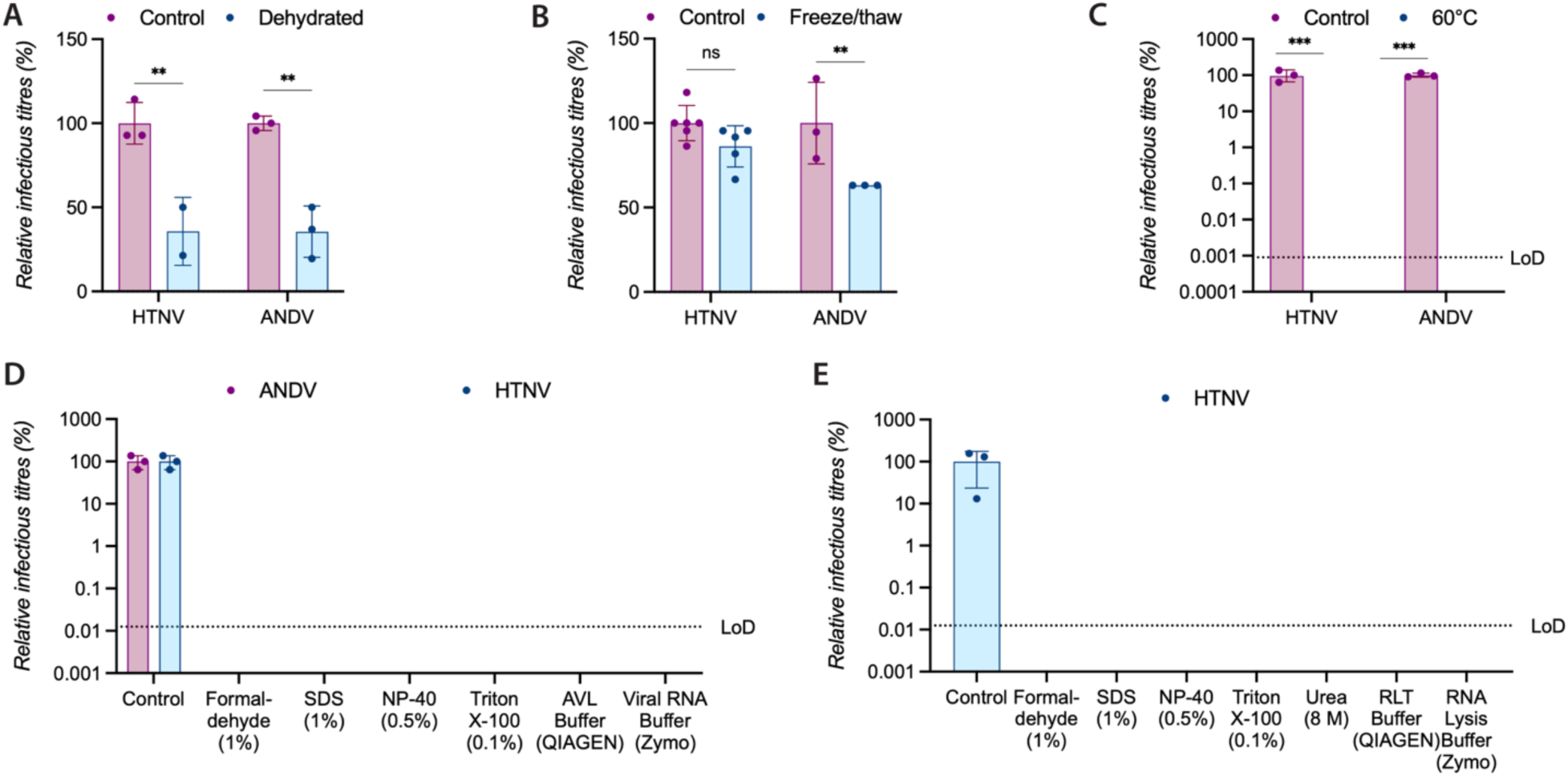
Validation of inactivation methods using highly pathogenic hantaviruses. (A) Relative infectious titres of HTNV or ANDV before and after dehydration at room temperature. (B) Relative infectious titres of HTNV or ANDV before and after a single freeze-thaw cycle. (C) Relative infectious titres of HTNV or ANDV before and after incubation for 5 min at 60°C. (D) Relative infectious titres of HTNV or ANDV before and after treatment with the indicated chemicals in solution. (E) Vero E6 cells were infected with HTNV for 2 dpi, when cells were inactivated with the indicated chemical reagents or a PBS control and ruptured by freeze-thawing. Cell lysates were used to infect naïve Vero E6 cells and infectious titres were determined at 20 dpi. All graphs represent relative data from immuno-plaque assays normalised to the control condition, with dots depicting independent biological replicates and error bars the standard deviation of the mean. The dotted lines represent the limit of detection (LoD) for each graph. Statistical significance was determined by two-way ANOVA followed by Šidák’s correction for multiple comparisons: *** (p > 0.001), ** (p > 0.01), * (p > 0.05), ns (p > 0.05).

Next, we performed chemical inactivation of cell-free HTNV and ANDV (Fig 5D) as previously shown for TULV. Analogous to TULV, we also observed a complete loss of infectious viral titres using 1% formaldehyde, 0.5% NP-40, AVL Buffer or Viral RNA Buffer for both HTNV (≥19,000-fold reduction over limit of detection) and ANDV (≥7,500-fold reduction over limit of detection). Notably, we also observed full inactivation using 1% SDS, 0.1% Triton X-100 and 8 M urea treatment, which in contrast only resulted in significant yet incomplete inactivation using TULV (Fig 3C).

Finally, we validated whether complete inactivation of HTNV infected cells could be achieved using fixation or commonly used cell lysis buffers. Here, we demonstrated that 20 days after transfer of cell lysates that had been treated with fixatives (1% formaldehyde), detergents (1% SDS, 0.5% NP-40, 0.1% Triton X-100, 8 M urea) or RNA extraction buffers (RLT Buffer, RNA Lysis Buffer) onto naïve cells, no infectious virus could be detected in the cell culture media (Fig 5E).

Overall, we were able to demonstrate that highly pathogenic hantaviruses such as ANDV and HTNV exhibit the same or even more susceptible behaviour as TULV when exposed to environmental stresses or inactivating chemicals and can be efficiently inactivated using commonly used laboratory conditions.

## Discussion

Emerging viruses have caused several global pandemics and local epidemic outbreaks in the past decades. Hantaviruses represent a particular threat to global health, due to their high prevalence in their natural host species, the ubiquitous abundance of their rodent host species and their close interactions with human populations, the highly pathogenic and lethal disease in humans as well as the proven potential for human-to-human transmission. Our need to study these viruses to understand basic molecular details as well as to develop effective and safe antivirals and vaccines is thus self-evident. Hantaviruses are primarily transmitted environmentally: viral particles are shed in bodily fluids and secretions such as rodent droppings, which might subsequently be accidentally ingested or inhaled by humans or other host species. Hantaviruses therefore rely on a certain virion stability to withstand environmental stresses in a cell-free context, such as dehydration or freezing of viral particles or prolonged exposure to high or low temperatures.

We demonstrated that when stored in cell culture media, TULV only loses a fraction (∼20%) of its infectivity after 21 days at 4°C, whereas at 37°C it rapidly loses its infectious potential with a half-life of 13 h, and the last infectious particles can be detected at 10 days (Fig 2D). These observations explain why the cumulative infectious viral titres in cell culture when growing TULV is not significantly different from the titres that were only produced in the last day or days (Fig 1B). These data are also consistent with a previous study, where it was demonstrated that HTNV can remain infectious for up to 7 days in cell culture media when stored at 37°C or for up to 9 days when stored at 21°C, but for up to 12 weeks when stored at 4°C [23]. However, another study showed that PUUV and TULV lost their infectivity after 24 h when stored at 37°C, after 11 days when stored at 23°C and still retained reduced infectivity after 18 days when stored at 4°C [24]. It is important to note though, that these previously published results for PUUV and TULV only were able to capture a very small dynamic range of virus titres (less than 50-fold reduction of infectivity), compared to several orders of magnitude (approximately 100,000-fold reduction of infectivity) that can be detected using immuno-plaque assays as used in the previously published HTNV study and in our own data. This might explain why infectious viral particles could be detected for longer periods of time by us using the same virus (TULV) compared to the previously published results. Interestingly, it was also demonstrated that PUUV remains infectious to naïve bank voles in bank vole droppings for up to 15 days [24]. While not equivalent to being fully solubilised in cell culture media, it is likely that the virus is at least partly protected from full dehydration and other stresses inside animal droppings.

As viral particles seldom remain suspended in liquid for prolonged times in the environment, we showed that dehydration of TULV, HTNV and ANDV particles results in a similar (∼65%) loss of infectivity (Fig 2E). These results are broadly concur with previous studies, where on the one hand, an approximate 10-fold reduction in infectious viral particles were observed after a few hours and no infectious virus remaining after 24 h for HTNV, PUUV and TULV [23,24]. And on the other hand, we also recently demonstrated that while ANDV particles undergo an approximately 1,000-fold reduction in infectivity after 24 h when dehydrated on steel discs, it can nevertheless remain infectious for more than 5 days, similar to our data for TULV which we could detect at low concentrations for up to 5 days [25]. Slight differences between the studies could perhaps be explain by different experimental setups, different methods for virus preparation and other environmental factors that are known to influence viral stability and that were not taken into account in these studies, such as humidity, light exposure, pH level and protein content of the virus stocks [26–30]

Many hantaviruses can only be handled in a high containment facility and need to be fully inactivated before they can be processed under normal biosafety levels. While inactivation methods have been extensively tested against a multitude of different viruses [31], there are not many studies which comprehensively and convincingly validate the safe inactivation of cell-free and cell-associated hantaviruses. We demonstrate that hantaviruses rapidly lose their infectivity after exposure to 56°C or higher and that after 30 seconds at 60°C infectious titres already are approximately 1,000-fold reduced and that no infectious particles remain after 5 min (Fig 2C-D and 5C). This confirms previous results that show that in solution PUUV is fully inactivated after 15 min at 56°C [24]. Interestingly, the same study also suggests that PUUV is able to withstand 56°C heat for up to 2 h when fully dehydrated, which appears to be in direct contradiction with with our own data demonstrating that TULV rapidly loses its infectivity even at 37°C (Fig 2F).

Recently, we already established that ANDV can effectively and quickly be inactivated using ethanol or 2-propanol-based WHO-recommended hand rub formulations [25], which was in line with previous studies using enveloped RNA and DNA viruses [26,32–34]. Furthermore, PUUV has been shown to be effectively inactivated using commercial disinfectants Clidox®, Halamid-d®, and Virkon® S [35]. In addition, cell-free HTNV was shown to be fully inactivated by 2 min exposure to 40% ethanol [23] and intracellular HTNV was fully inactivated using 100% methanol (8 min), 1% paraformaldehyde (20 min) or 1:1 acetone/methanol (10 min) [36]. Our own results confirm that full inactivation of hantaviruses can be achieved using 40% ethanol (cell-free virus) or 1% formaldehyde (cell-free and cell-associated virus) (Fig 3A-B, 4A, 5D-E). In addition, we also provide evidence for the complete inactivation of hantaviruses using different commercial RNA extraction kits (Fig 3D, 4C, 5D-E) and different detergents – including SDS, NP-40, Triton X-100 and urea -used in common lysis buffers (Fig 3C, 4B, 5D-E). Interestingly, our results suggest that ANDV and HTNV are slightly more susceptible to SDS and Triton X-100 than TULV, which exhibited partial resistance to complete inactivation using 1% SDS or 0.1% Triton X-100 alone. While more studies would be needed to investigate these differences, it is also important to highlight that we only tested single reagents in isolation and that commonly combinations of different inactivation methods are used, *e.g.* several inactivating reagents in lysis buffers or a combination of lysis buffers and heat inactivation.

Overall, these data are broadly in line with similar studies validating inactivation methods for other highly pathogenic RNA viruses, including arenaviruses and Nipah virus [37–39]. However, our study often establishes a lower necessary baseline for inactivation compared to many traditional inactivation methods, indicating that highly pathogenic viruses often are inactivated at much lower concentrations or exposure times than commonly used in the laboratory.

Together these data establish safe and effective inactivation methods compatible with common downstream molecular biology, cell biology, immunology or virology assays, including nucleic acid extraction, western blots, co-immunoprecipitation assays, ELISA, immunofluorescence assays and flow cytometry. Crucially, we demonstrate that our findings are not only applicable to a single hantavirus species, but we that are broadly valid across a diverse set of Old World and New World hantaviruses, including the highly pathogenic HTNV and ANDV. Moreover, our data also goes a long way to detail the environmental stability of hantavirus particles with strong implications for our understanding of the environmental transmission potential of hantaviruses.

## Acknowledgements

A.A. was supported by the Hannover Biomedical Research School (HBRS) and the Center for Infection Biology (ZIB). This study was also funded by the Deutsche Forschungsgemeinschaft (DFG, German Research Foundation) under Germany’s Excellence Strategy -EXC 2155 -project number 390874280 (to B.E.N. and T.P.) and the Swedish Research Council (2023-02595 and 2024-03783 to B.E.N.).

